# Metronomic capecitabine as an immune modulator in glioblastoma patients reduces myeloid-derived suppressor cells

**DOI:** 10.1101/655688

**Authors:** David M. Peereboom, Tyler J. Alban, Matthew M. Grabowski, Alvaro G. Alvarado, Balint Otvos, Defne Bayik, Gustavo Roversi, Mary McGraw, Pengjing Huang, Alireza M. Mohammadi, Harley I. Kornblum, Manmeet S. Ahluwalia, Michael A. Vogelbaum, Justin D. Lathia

**Affiliations:** Rose Ella Burkhardt Brain Tumor and Neuro-Oncology Center, Cleveland Clinic, Cleveland, Ohio, 44195, USA; Case Comprehensive Cancer Center, Case Western Reserve University, Cleveland, Ohio, 44106, USA; Cancer Impact Area and Department of Cardiovascular & Metabolic Sciences, Lerner Research Institute, Cleveland Clinic, Cleveland, Ohio, 44195, USA; Department of Molecular Medicine, Cleveland Clinic Lerner College of Medicine at Case, Western Reserve University, Cleveland, Ohio, 44195, USA; Department of Psychiatry and Biobehavioral Sciences and Semel Institute for Neuroscience, University of California Los Angeles, Los Angeles, California, 90095, USA; Department of NeuroOncology, Moffitt Cancer Center, Tampa, Florida, 33612, USA

## Abstract

**Background:** Myeloid-derived suppressor cells (MDSCs) are elevated in glioblastoma (GBM) patient circulation, present in tumor tissue, and associated with poor prognosis. While low-dose chemotherapy reduces MDSCs in preclinical models, the use of this strategy to reduce MDSCs in GBM patients has yet to be evaluated.

**Methods:** A phase 0/1 dose-escalation clinical trial was conducted in recurrent glioblastoma patients treated 5-7 days prior to surgery with low-dose chemotherapy via capecitabine followed by concomitant low-dose capecitabine and bevacizumab. Clinical outcomes, including progression-free and overall survival, were measured, along with safety and toxicity profiles. Over the treatment time course, circulating MDSC levels were measured by multi-parameter flow cytometry, and tumor tissue immune profiles were assessed via mass cytometry time-of-flight.

**Results:** A total of 11 patients were enrolled across escalating dose cohorts of 150, 300, and 450 mg bid, with a progression-free survival of 5.8 months (range of 1.8-27.8 months) and an overall survival of 11.5 months (range of 3->28.0 months) from trial enrollment. No serious adverse events related to the drug combination were observed. Compared to pre-treatment baseline, circulating MDSCs were found to be higher after surgery in the 150 mg treatment arm and lower in the 300 mg and 450 mg treatment arms. Increased cytotoxic immune infiltration was observed after low-dose capecitabine compared to untreated GBM patients in the 300 mg and 450 mg treatment arms.

**Conclusions:** Low-dose, metronomic capecitabine in combination with bevacizumab is well tolerated in GBM patients and was associated with a reduction in circulating MDSC levels and an increase in cytotoxic immune infiltration into the tumor microenvironment.

**Trial registration:** NCT02669173

**Funding:** This research was funded by the Cleveland Clinic, Case Comprehensive Cancer Center, Musella Foundation, and B*CURED. Capecitabine was provided in kind by Mylan Pharmaceuticals.

## Introduction

Glioblastoma (GBM) is the most common primary malignant brain tumor, with an annual incidence of 11,000 cases in the U.S. and a median survival of 14-18 months despite aggressive therapies^1–5^. Currently, standard of care includes maximal safe surgical resection followed by concomitant radiation and chemotherapy with temozolomide^1^. In nearly 100% of cases, this approach fails, resulting in a recurrent tumor with limited therapeutic options. Few therapies are FDA-approved for patients with recurrent GBM, including lomustine, bevacizumab, carmustine wafers, and tumor-treating fields, none of which have demonstrated a marked improvement in overall survival^6–9^. Multiple other cancers have faced similar obstacles to effective treatment, but this has been recently overcome with the use of immune-modulating therapies, and as a consequence, there is interest in trying to modify the immune system in GBM. Several immunomodulatory approaches are currently under clinical evaluation, including the use of immune checkpoint inhibitors, oncolytic viruses, dendritic cell vaccines, and CAR-T cell approaches^10^, but the well-appreciated immunosuppressive nature of GBM has proven difficult to overcome^11–15^.

A hallmark of GBM immunosuppression is the appearance of circulating myeloid-derived suppressor cells (MDSCs) at higher levels than in many other cancers^11, 13, 16–20^. This heterogeneous cell population is activated upon injury and in many cancers, where MDSCs inhibit cytotoxic immune cell populations and contribute to overall immune suppression^21–25^. In multiple solid tumor models and clinical trials, elevated peripheral MDSC levels have been correlated with a more immunosuppressive phenotype, as well as with tumors that were refractory to immune-activating therapies, including immune checkpoint inhibitors^26, 27^. We previously observed that GBM patients with a better prognosis had reduced MDSCs in their tumors as well as in their peripheral circulation^13, 22^. Previous studies have demonstrated that MDSCs in multiple tumor types can be reduced via low-dose chemotherapies^28–30^. We recently found that this could be achieved in pre-clinical GBM mouse models via 5-fluorouracil (5-FU), an antimetabolite drug that impacts both RNA and DNA synthesis via its inhibition of thymidylate synthase. Utilizing a metronomic low-dose 5-FU strategy, we were able to reduce circulating MDSCs, increase intratumoral activated T cell populations, and prolong survival^11^. Based on these pre-clinical observations, we sought to test this approach in recurrent GBM patients using an orally bioavailable 5-FU prodrug, capecitabine, combined with standard-of-care bevacizumab, an anti-angiogenic agent. The addition of bevacizumab, an anti-VEGF (vascular endothelial growth factor) antibody, was used to ensure that patients received standard therapy for recurrent GBM. In this phase 0/1 dose-escalation trial, our objective was to assess the ability of capecitabine to reduce circulating and tumoral MDSCs and to determine the safety/toxicity profile of capecitabine alone and in combination with bevacizumab in this patient population. Based on the hypothesis that this treatment approach would reduce immune suppression, we also analyzed circulating immune cells via flow cytometry and evaluated the immune profile of treated tumors with mass cytometry time of flight (CyTOF).

## Results

### Phase 0/1 clinical trial demographics

Patient accrual began in October 2016. To target MDSCs, we assessed the efficacy of escalating low doses of metronomic capecitabine with a fixed, standard dose of bevacizumab in a clinical trial approved by the Cleveland Clinic IRB (NCT02669173). Once GBM recurrence was identified on MRI, study-specific informed consent was obtained. Capecitabine was given 5-7 days prior to operation and then continued in 28-day cycles with periodic blood draws to assess peripheral blood immune cell populations over the course of the trial (Figure 1). A total of 12 patients were enrolled initially, with one patient removed from all aspects of the study after their resection specimen identified pseudoprogression. The demographics of the 11 evaluable patients are summarized in Table 1. The median age at diagnosis was 58 years; 7 patients were enrolled at the time of their first progression, and the remaining 4 patients were enrolled at their second progression. Surgical therapy at diagnosis included biopsy only (2 patients), biopsy with laser ablation (1 patient), and surgical resection (8 patients). All patients then received standard-of-care radiation with concurrent and adjuvant temozolomide. Additional therapies prior to trial enrollment included other chemotherapies (4 patients (2 had received lomustine and 2 received tesevatinib a, a multi-targeted tyrosine kinase inhibitor)), tumor-treating fields (1 patient), and immunotherapies (3 patients). The 3 patients who previously received immunotherapy included 2 patients who failed anti-LAG3 treatment and 1 patient who received the SurVaxM vaccine. Five patients had MGMT-methylated GBMs and 1 of these had an IDH mutation. These 3 patients demonstrated no remarkable differences in immune populations compared to the others on the trial. Expanded patient details, including molecular markers and other disease-management paradigms, are provided in **Supplemental Table 1**. Patients received capecitabine in 3 dose cohorts: 150 mg bid (4 patients), 300 mg bid (3 patients), and 450 mg bid (4 patients).

**Table 1.** Demographic, clinical, and tumor characteristics of the patients enrolled in the trial. Molecular markers for each patient were not available for every test; percentages are shown out of the total patients who had testing done for that specific marker.

| Variable | Median Value (range) |
| --- | --- |
| Total Patients | 11 |
| Age at Diagnosis | 58 (38-67) |
| Male Sex | 6/11 (55%) |
| Karnofsky Performance Status | 90 (30-100) |
| 90 | 7/11 (64%) |
| 80 | 4/11 (36%) |
| Ethnicity |  |
| White | 11/11 (100%) |
| Molecular Markers |  |
| 1p Loss | 2/9 (22%) |
| 19q Loss | 2/9 (22%) |
| IDH Mutated | 1/11 (9%) |
| MGMT Methylated | 5/11 (45%) |
| EGFR Amplification | 5/9 (56%) |
| Initial Surgical Therapy |  |
| Biopsy only | 2/11 (18%) |
| NTR | 3/11 (27%) |
| GTR | 5/11 (45%) |
| Laser Ablation | 1/11 (9%) |
| Prior Local Therapy |  |
| External beam radiation therapy | 11/11 (100%) |
| Tumor Treating Fields | 1/11 (9%) |
| Prior Systemic Therapy |  |
| TMZ | 11/11 (100%) |
| Immunotherapy | 3/11 (27%) |
| Chemotherapy | 4/11 (36%) |
| Subsequent Chemotherapy | 3/11 (27%) |
| Progression Number at Trial Enrollment |  |
| First | 7/11 (64%) |
| Second | 4/11 (36%) |
| Extent of resection |  |
| STR | 2/11 (18%) |
| NTR | 5/11 (45%) |
| GTR | 4/11 (36%) |
| Study Dose |  |
| 150 mg bid | 4/11 (36%) |
| 300 mg bid | 3/11 (27%) |
| 450 mg bid | 4/11 (36%) |
| Number of Cycles Completed |  |
| 150 mg bid* | 7.5 cycles (4-30) |
| 300 mg bid | 5 cycles (2-6) |
| 450 mg bid* | 8 cycles (2-14) |
| Overall | 6 cycles (2-30) |
| Progression Free Survival at 6 months | 5/11 (45%) |
| Progression Free Survival | 5.8 months (1.8-27.8) |
| Overall Survival | 11.5 months (3.0-28.0) |
| Number of Patients Still Active on Trial at Publication** | 2/11 (18%) |
| Number of Patients Still Alive at Publication ** | 3/11 (27%) |

Abbreviations: IDH: isocitrate dehydrogenase; MGMT: O-6-methylguanine-DNA methyltransferase; EGFR: epidermal growth factor receptor; STR: subtotal resection; NTR: near-total resection; GTR: gross-total resection; XRT: external beam radiation therapy; TMZ: temozolomide; bid: twice daily
\* One patient in each group is still active on trial
\*\* As of 05/01/2019

**Figure 1.**
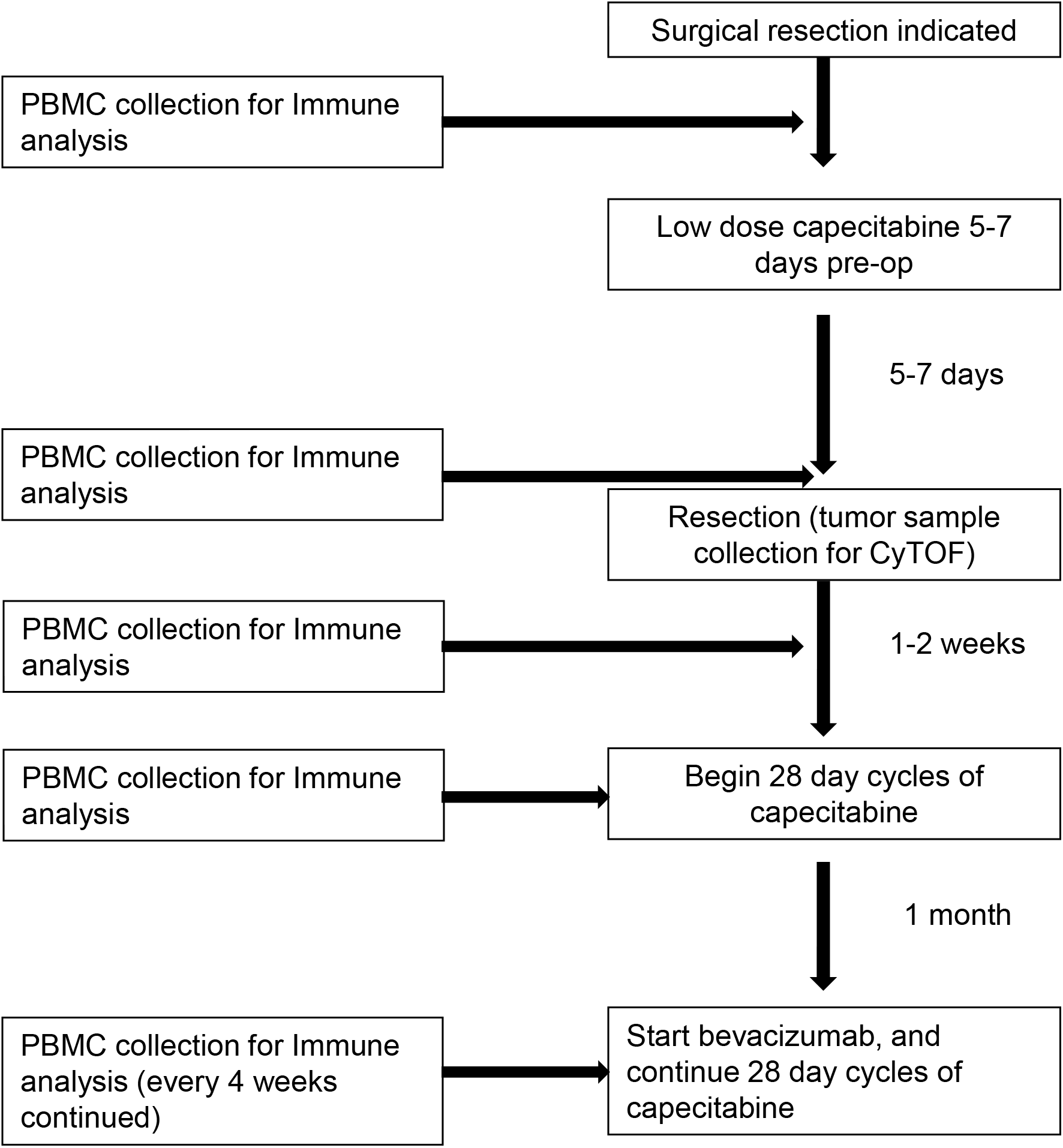
Study schematic demonstrating the time points for capecitabine treatment and immune analysis via flow cytometry and CyTOF.

### Clinical outcomes

The median follow-up time was 10.7 months (range 3 to 29 months). The median progression-free survival (PFS) was 5.8 months (ranging from 1.8 months to >27 months, Figure 2A), with 45% (5 out of 11 patients) achieving a 6-month PFS. The median overall survival (OS) was 11.5 months from trial enrollment (ranging from 3.0 to >28.0 months, Figure 2B). The number of patients still active on trial at the time of manuscript submission is 2 out of 11 (18%), and the number of patients alive is 3 out of 11 (27%). The median follow-up time was 10.7 months with a range of 3-29 months. The median PFS was 7.25 months for patients receiving capecitabine at 150 mg bid, 5.5 months for those receiving 300 mg bid, and 7.35 months for those receiving 450 mg bid. The median OS was 16.6 months for patients receiving capecitabine at 150 mg bid, with one patient still on study; 11.5 months at 300 mg bid, and 9.8 months at 450 mg bid, with one patient still on the study and 1 patient alive but taken off the study due to progressive disease.

**Figure 2.**
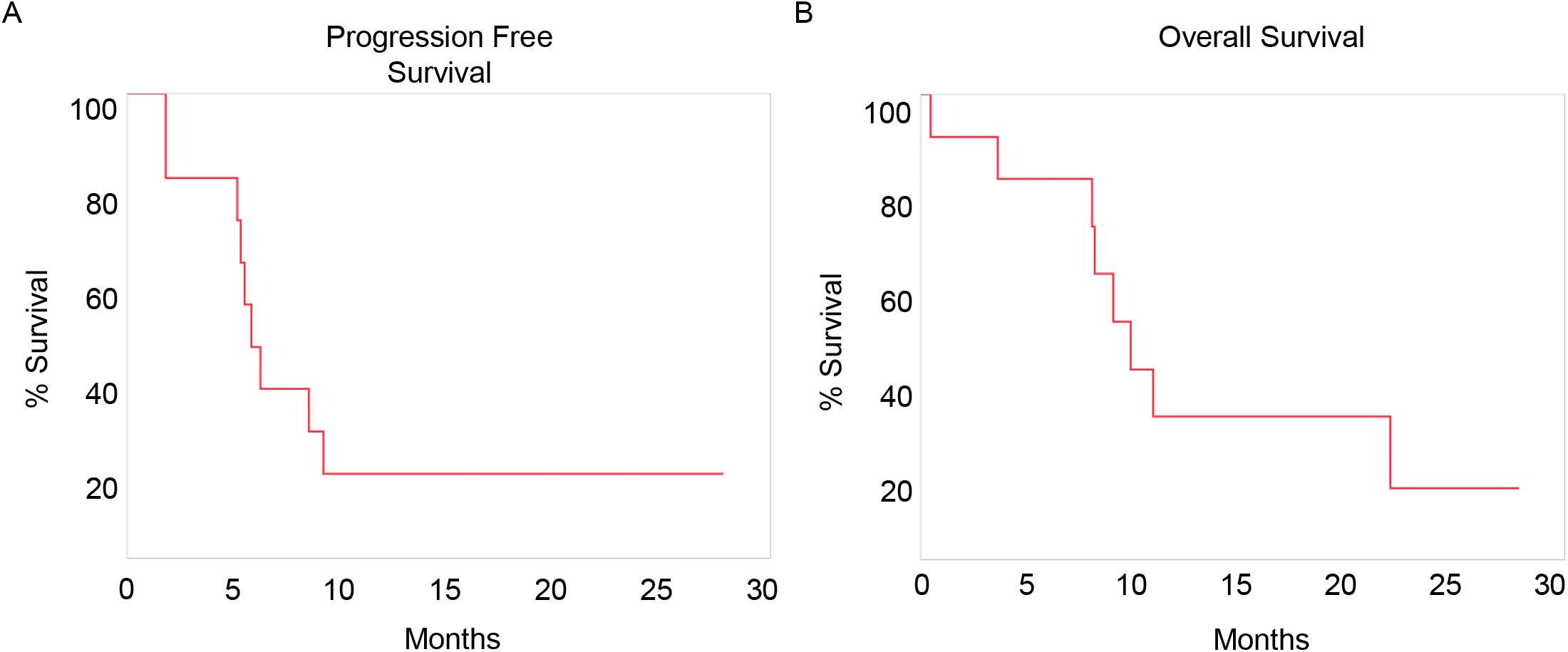
Kaplan-Meier analysis of progression free survival and overall survival. Progression free survival (**A**) and overall survival (**B**) of enrolled patients are represented as Kaplan Meier survival curves.

### Tolerability of metronomic low-dose capecitabine therapy in recurrent GBM patients

Capecitabine at all three doses administered in combination with bevacizumab was generally well tolerated, with one treatment-related serious adverse effect (event possibly, probably, or definitely related to treatment) – grade 5 perforated diverticulum (Table 2). This patient suffered the event after going off study but within 30 days of completion of the study treatment. No other grade 4 or 5 events occurred. Tables 2 and 3 summarize the treatment-related adverse events, which included one patient each with grade 3 thromboembolism and dyspnea. One patient in the 450 mg bid cohort experienced grade 3 anemia approximately 2.5 months into the trial and was dose-reduced to 300 mg bid. Other side effects were grade 1 and 2 and included fatigue (4 patients), hypertension (3 patients), nausea and vomiting (2 patients), and a small intracranial hemorrhage that required no intervention (1 patient).

**Table 2.** Causes of each patient being removed from the study, grouped by capecitabine dose. There are two patients still on the trial receiving treatment.

| **Reason Off Study (9 patients\*)** | **No. (%)** |
| --- | --- |
| Dose 1 – 150 mg bid – 4 patients |  |
| Radiographic/Clinical Progression | 2/4 (50%) |
| Adverse Event/Side Effects/Complications | 1/4 (25%) |
| Still receiving Study Treatment as of 5/1/2019 | 1/4 (25%) |
| Dose 2 – 300 mg bid – 3 patients |  |
| Radiographic/Clinical Progression | 3/3 100%) |
| Dose 3 – 450 mg bid – 4 patients |  |
| Radiographic/Clinical Progression | 2/4 (50%) |
| Patient Withdrawal for Other Reasons | 1/4 (25%) |
| Still receiving Study Treatment as of 5/1/2019 | 1/4 (25%) |

\*2 additional patients receiving treatment as of 5/1/19

**Table 3.** Treatment-related adverse events that were graded as possibly, probably, or definitely related to the treatment (capecitabine + bevacizumab). There were no Grade 4 or 5 events.

| Treatment-Related Adverse Events (n =11 patients) | No. of Events by Grade |  |  |  | No. of Patients |
| --- | --- | --- | --- | --- | --- |
|  | Grade 1 | Grade 2 | Grade 3 | Grade 5 |  |
| Hematological Adverse Events |  |  |  |  |  |
| Anemia |  |  | 1 |  | 1/11 (9%) |
| Thromboembolic Event |  |  | 1 |  | 1/11 (9%) |
| Thrombocytopenia |  | 1 |  |  | 1/11 (9%) |
| Easy Bruising | 1 |  |  |  | 1/11 (9%) |
| Nervous System Adverse Events |  |  |  |  |  |
| Intracranial Hemorrhage |  | 1 |  |  | 1/11 (9%) |
| Peripheral Sensory Neuropathy | 1 |  |  |  | 1/11 (9%) |
| Gastrointestinal Adverse Events |  |  |  |  |  |
| Nausea/Vomiting | 1 | 1 |  |  | 2/11 (18%) |
| Constipation | 2 |  |  |  | 2/11 (18%) |
| Perforated Diverticulum |  |  |  | 1* | 1/11 (9%) |
| Abdominal Pain | 1 |  |  |  | 1/11 (9%) |
| Oral Mucositis | 1 |  |  |  | 1/11 (9%) |
| Early Satiety | 1 |  |  |  | 1/11 (9%) |
| Other Adverse Events |  |  |  |  |  |
| Fatigue | 2 | 2 |  |  | 4/11 (36%) |
| Hypertension | 2 | 1 |  |  | 3/11 (27%) |
| Dyspnea | 1 |  | 1 |  | 2/11 (18%) |
| Fever | 1 |  |  |  | 1/11 (9%) |
| Arthralgia | 1 |  |  |  | 1/11 (9%) |

### Daily oral capecitabine reduced the expected increase in MDSCs post-surgical resection

Peripheral blood MDSC concentrations were evaluated via flow cytometry (**Supplemental Figure 1A**) before treatment, during surgical resection and post-surgical resection on a per-patient basis to visualize trends in MDSC changes over time^13^ (Figure 3A). Patients in the 300 mg bid and 450 mg bid capecitabine treatment arms had a reduction in MDSCs post-surgery, which was not observed in the 150 mg bid arm (Figure 3A). As a comparison, a longitudinal analysis of newly diagnosed GBM patients conducted in a separate, contemporaneous study showed a continued increase in MDSCs over time after surgical resection (Figure 3A). This increase in peripheral blood MDSCs post-surgery has been attributed to surgical intervention in multiple cancer types^16, 23, 31, 32^ and occurs in untreated patients prior to surgical resection (Figure 3A). These analyses allowed us to identify a relative reduction in peripheral MDSCs post-surgical resection in the 300 and 450 mg bid capecitabine cohorts in 6 out of 7 patients (Figure 3A). Of note, patient 3 in the 450 mg bid capecitabine treatment cohort (the only patient who did not have a reduction in peripheral MDSCs post-surgery) was noted to have multifocal GBM at the time of recurrence and enrollment in the trial. The distal site of recurrence was not resected, and the patient progressed 1 month later at that lesion site. Therefore, this patient was excluded from additional post-surgical analysis of peripheral blood immune populations. A comparison of the average fold change in MDSCs post-surgical resection demonstrated an increase of MDSCs in untreated patients, a return to pre-operative baseline in patients treated with 150 mg bid capecitabine, and a reduction in patients treated with both 300 mg bid and 450 mg bid capecitabine. The 300 mg bid capecitabine treatment reduced MDSCs to a level that was not further reduced when the capecitabine treatment was increased to 450 mg bid (Figure 3B). Using a similar flow cytometry approach, analysis of peripheral T cell populations (CD3+, CD4+, CD8+, T regulatory cells) showed no change in the circulation at any dose of capecitabine or in response to surgery (**Supplemental Figure 1B-F**).

**Figure 3.**
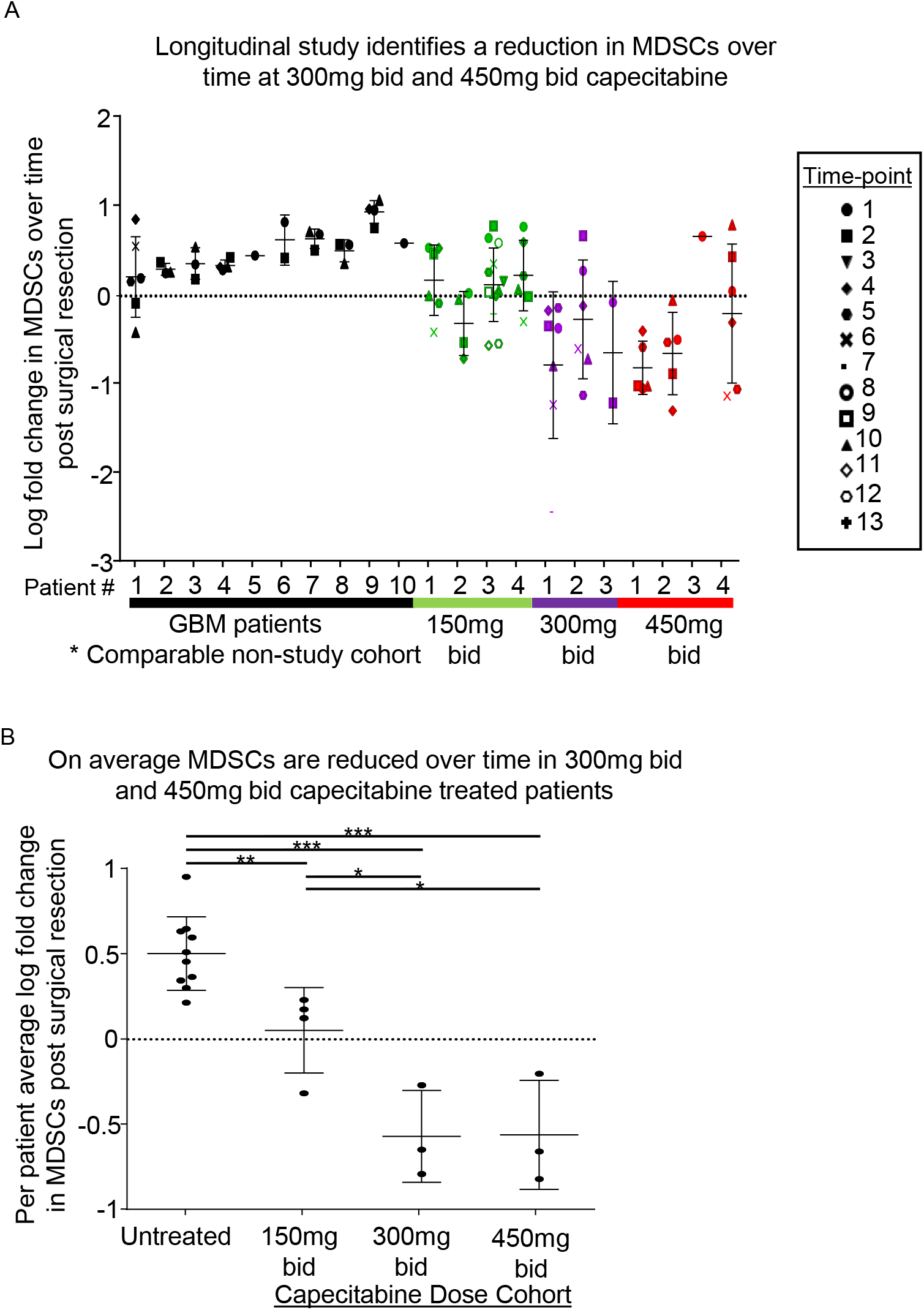
Peripheral MDSCs are reduced over time in patients treated with capecitabine at 300 mg bid and 450 mg bid. Flow cytometry analysis of PBMCs longitudinally identified MDSCs (HLA-DR^−/low^, CD11b^+^, CD33^+^), and the log fold change in MDSCs per patient post-surgical resection is depicted (**A**), with each symbol representing the blood draws in sequential order from 1-13. The average log fold change of MDSCs per patient over time is graphed per treatment group (**B**) and identified a significant difference between untreated and all treatment groups and a maximal reduction in the 300 mg bid and 450 mg bid treatment groups (**B**). All error bars represent the standard deviation. Unpaired student’s t-test was used for all comparisons where *p < 0.05, ** p < 0.01, ***p < 0.001.

To further determine the changes in tumor immune profiles associated with systemic capecitabine treatment, we analyzed GBM tissue from patients treated with capecitabine 5-7 days before surgery via CyTOF, which we previously used to identify immune shifts associated with GBM patient prognosis^13^. This immune panel consisted of 28 key cell surface immune system markers (**Supplemental Figure 2A**). In the CD45+ cell fraction from cryopreserved single-cell tumor suspensions of newly diagnosed GBM patients, recurrent GBM patients, and GBM patients treated with capecitabine in our trial (300 mg and 450 mg bid, Figure 4A), we identified 29 unique immune populations in an unbiased manner by using t-distributed stochastic neighbor embedding (t-SNE) analyses (Figure 4B, Supplemental Figure 2B, 2C) with no major change in overall CD45+ cell numbers (**Supplemental Figure 2B**). When comparing patients with newly diagnosed GBM, recurrent GBM, and recurrent GBM from the capecitabine-dose cohorts, we observed shifts in the tumor-infiltrating immune cell population (Figure 4C, **Supplemental Figure 3**). Overall, the newly diagnosed and recurrent patients appeared to have similar populations of immune cell clusters, while the groups treated with 300 mg bid and 450 mg capecitabine demonstrated a distinct immune phenotype resembling a more immune activated status (Figure 4C, **Supplemental Figure 3**). We did not observe a substantial change in overall intratumoral lymphocyte number after capecitabine treatment (**Supplemental Figure 2B**). When comparing these patients to untreated patients with GBM (including newly diagnosed and recurrent), we observed significant increases in CD4+ central memory T cells (subset 1), CD8+ effector memory cells, classical monocytes, dendritic cells, macrophages, microglia (subset 1), NK cells (CD56 high), and NK cells (CD56 mid) (Figure 5, **Supplemental Figure 4A, B**). In addition, the application of a recently published machine-learning algorithm using the R package CytoDx^33^ to the intratumoral CyTOF data revealed that cytotoxic T lymphocyte-associated antigen 4 (CTLA-4) levels were the most predictive marker to distinguish untreated versus capecitabine-treated patients, and this is represented as a decision tree of predictions showing the predicted cell population changes that occur upon treatment (Figure 6A). Multidimensional (t-SNE) analyses of untreated vs capecitabine-treated tumor lymphocytes identified a reduction of CTLA-4 expression (Figure 6B). Manual gating of CyTOF data validated this prediction and identified a significant reduction in CTLA-4 expression in lymphocytes in capecitabine-treated patients (Figure 6C) and also confirmed a significant reduction in CTLA-4+/PD-1+ (programed cell death protein 1) macrophages (Figure 6D). Taken together, these data demonstrate that low-dose systemic capecitabine treatment reduces circulating MDSCs when compared to a GBM resected under standard conditions, with the latter group experiencing a surgery-induced spike. In addition, low-dose capecitabine altered the tumor immune microenvironment, enhancing the number and immunophenotype of cells associated with an anti-tumor response.

**Figure 4.**
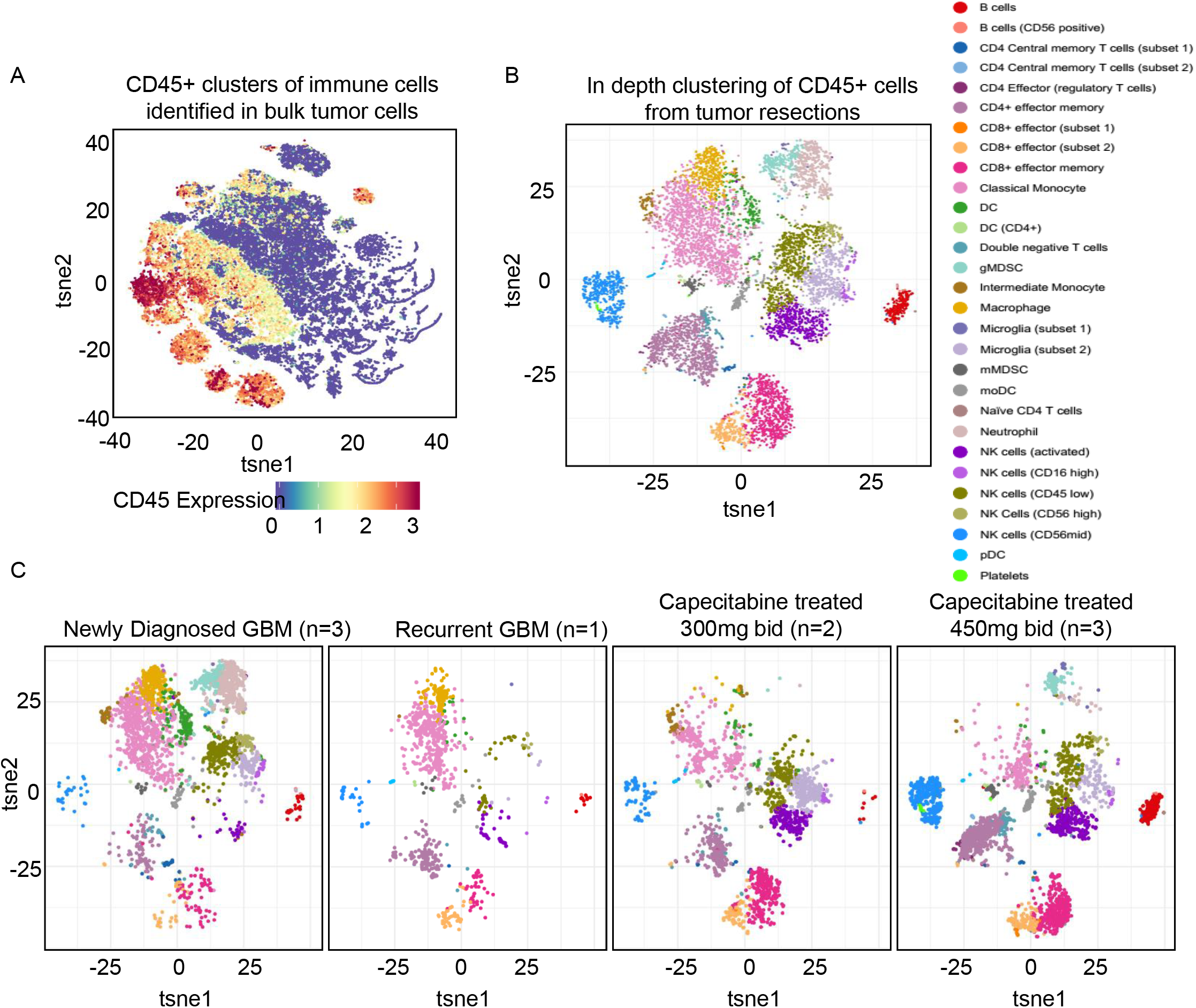
Capecitabine increases the immune activation in tumors after 7 days of treatment prior to surgery. CyTOF analysis using an immune panel of 28 immune markers analyzed capecitabine-treated tumor samples from patients 5, 6, 9, 10, and 11 along with newly diagnosed tumor samples (ND) (patients 1, 2, and 3), and one recurrent (Rec) GBM tumor (sample 1) (**A**) is represented as a tSNE multi-dimensional plot and colored by CD45 expression, highlighting the immune populations. After selecting immune populations based on CD45 expression, all tumor sample immune cells were combined and used to cluster immune populations in an unbiased manner from Live/CD45+ cells only (**B**). Separate newly diagnosed GBM patients (n=3), recurrent GBM patients (n=1), 300 mg bid capecitabine-treated patient (n=2), and 450 mg bid capecitabine-treated patients (n=3) tSNE plots represent the immune landscape of each tumor cohort (**C**).

**Figure 5.**
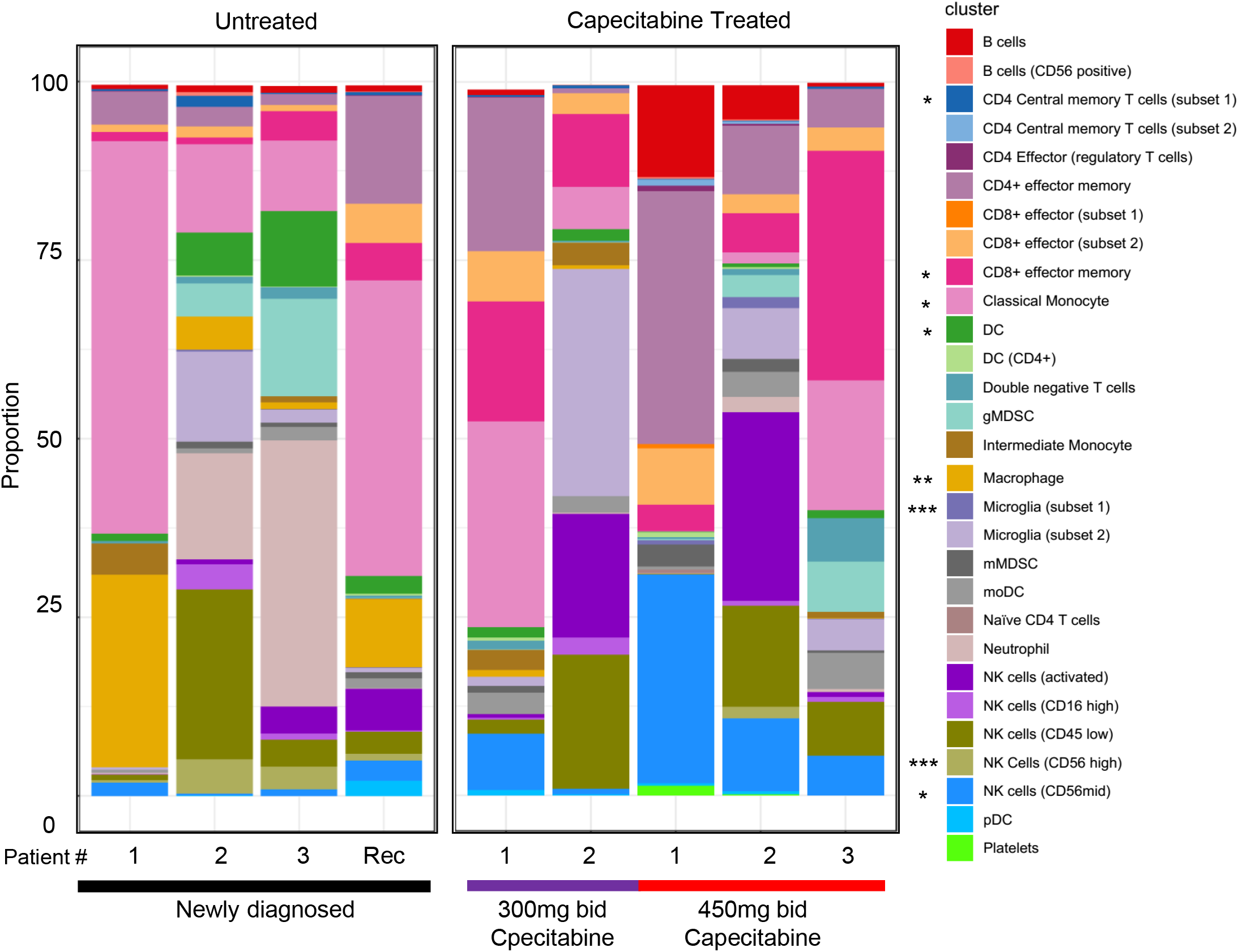
Comparison of untreated vs capecitabine-treated immune populations on a per-patient basis. Unbiased clustering identification of immune populations and quantification of the proportion of each cell type present in the CD45+ population is represented as a proportion of the total Live/CD45+. Statistical analysis comparing untreated vs treated identified statistically significant differences between the populations with an asterisk (*) to the left of the population color. Linear models of the data with two-tailed t-test comparisons and Benjamini-Hochberg were used to control for multiple comparisons. *p < 0.05, ** p < 0.01, ***p < 0.001. Graphs represent data sets as median with first and third quartiles.

**Figure 6.**
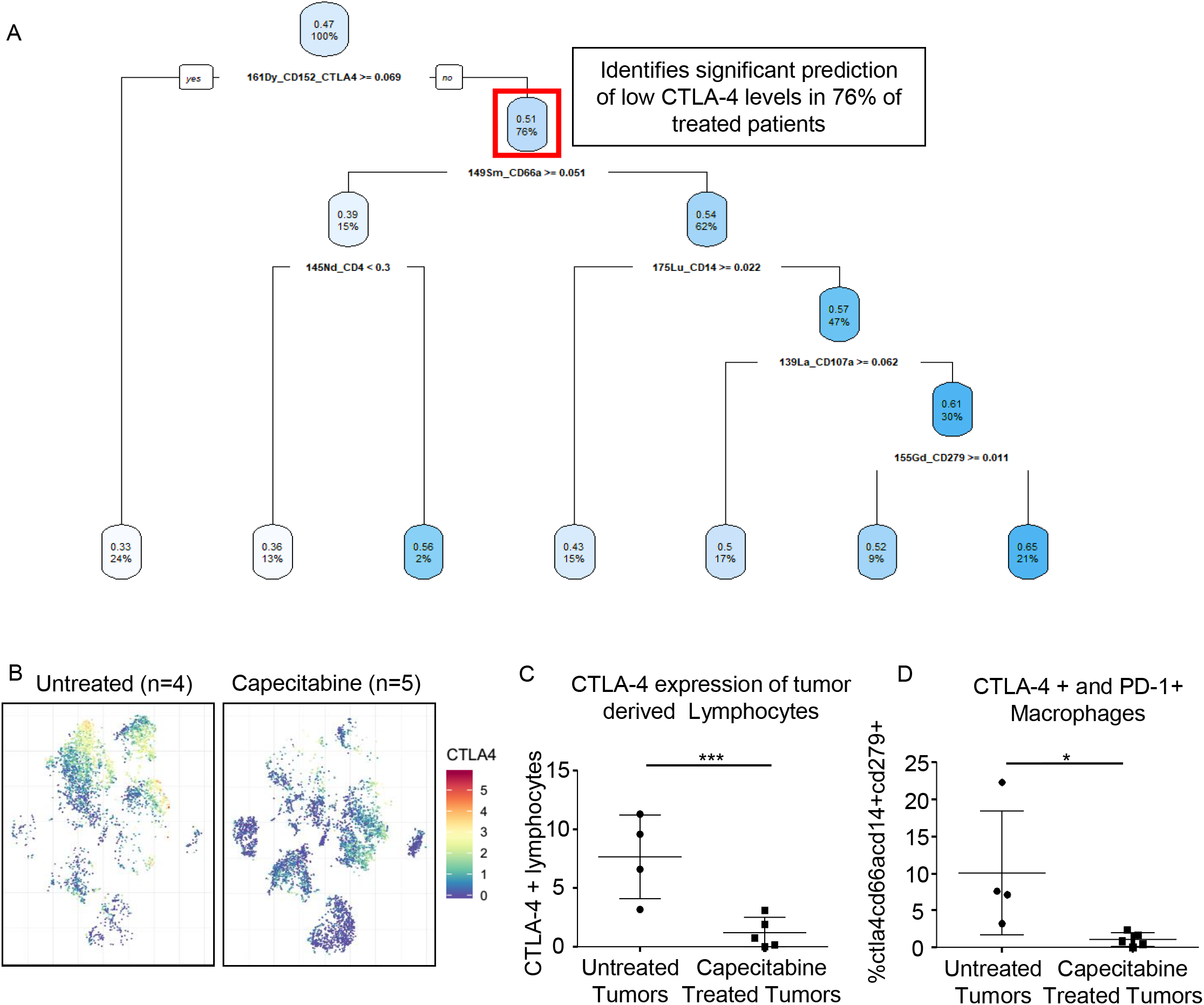
Utilizing patient tumor CyTOF data, a machine-learning approach identified a reduction in a signature for immune cell exhaustion in the tumors of capecitabine-treated patients. From the CyTOF data, a decision tree was generated using the CytoDx R package (**A**). The first node of the decision tree is highlighted, identifying the initial finding of 76% of patients with a lower level of CTLA-4+ cells. Multidimensional tSNE modeling of the total CD45+ cells from the tumors of untreated and treated patients colored by CTLA-4 expression levels identifies the clusters with a reduction in CTLA-4 upon treatment (**B**). Manual gating of the CyTOF data highlighted the quantitative differences in CTLA-4+ lymphocytes in the tumors of patients treated with capecitabine (**C**). Further manual gating for the final subset of CTLA-4-cells identified by the decision tree revealed a unique population of CTLA-4+, PD-1+ macrophages that were suppressed upon capecitabine treatment (**D**).

## Discussion

Findings from this study indicate that low-dose capecitabine pre-operatively and post-operatively in combination with bevacizumab is well tolerated at all doses, with an acceptable side-effect profile. While there is no control arm in this phase 0/1 trial, preliminary findings indicate that PFS and OS were comparable to historical controls^1, 7^. Comparisons of the immune populations between the capecitabine-treated groups revealed that 300 mg bid was the optimal capecitabine dose that led to maximal decreases in circulating MDSCs in a subset of patients without impacting lymphoid populations needed for an anti-tumor immune response. Finally, CyTOF analyses revealed that 5-7 days of priming the immune system with low-dose capecitabine treatment at 300 mg bid and 450 mg bid enhanced the anti-tumor immune cell populations within the tumor, including CD8^+^ effector memory cells and NK cells. However, these results are underpowered to resolve the complete effects of capecitabine on the tumor immune responses of the patients and support the expansion of this strategy to a larger, controlled clinical trial utilizing 300 mg bid capecitabine.

While patients on this trial received both capecitabine and bevacizumab, the tumor immune profile was analyzed on surgical samples obtained before bevacizumab treatment. Therefore, the observed intratumoral effects of capecitabine could not have been a result of bevacizumab treatment. Furthermore, although circulating MDSC levels were analyzed while patients were on bevacizumab and capecitabine, the noted MDSC changes began to occur before the initiation of bevacizumab. The decreasing MDSC levels also occurred in a capecitabine dose-dependent manner, making it unlikely that bevacizumab played a significant role. However, the potential adjuvant effects of bevacizumab on circulating immune populations cannot be inferred from these results.

In this trial, we employed an emerging single-cell phenotyping approach, CyTOF, to gain insight into comprehensive immune signatures within GBM. By integrating these data with a machine-learning approach^33^, we identified a potential increase in immune activation after capecitabine treatment. Recently, utilization of next-generation technology to enhance clinical trials identified key changes between responders and non-responders^34–36^. For example, in three recent GBM immunotherapy clinical trials, single-cell RNA-sequencing, CyTOF, and T cell receptor repertoire/clonality analyses demonstrated changes in the immune microenvironment as a result of therapy, immune alterations as a result of the dynamics of clonal tumor cell evolution, and specific immune signatures associated with response^34–36^. Taken together, these approaches demonstrate insight that can be provided by these immune-monitoring approaches.

The dosing of chemotherapy agents has historically focused on the maximal tolerated dose to increase anti-tumor effects. In many cases, these high doses are associated with profound bone marrow toxicity, eliminating both pro- and anti-tumor immune cells^28, 37–40^. However, chemotherapies such as 5-fluorouracil and gemcitabine differentially impact the immune system at low doses, and this has recently been observed with temozolomide^12, 16, 29, 41^. By reducing the dose of these standard chemotherapies, it may be possible to reduce a subset of immune cells that drive immune suppression with little negative impact on anti-tumor immune cell concentration or function. While the mechanisms of action are not well elucidated, this effect may be due to differential sensitivity and/or different proliferation rates. In this trial, the capecitabine dose (300 mg bid) that led to the maximal decrease in peripheral immunosuppressive MDSCs was four- to five-fold lower than that used in colorectal cancer (1250 mg bid)^42^ and pancreatic cancer (2000 mg bid) ^43^, and increasing the dose of drug (to 450 mg bid) did not offer further reductions in immunosuppressive MDSCs. Importantly, these lower doses of capecitabine increased intratumoral immune-activating cell subsets, suggesting that these doses were more specific for immunosuppressive compartments. Previously our preclinical models have illustrated this strategy of selective MDSC inhibition^11^, and with the results of this trial, we have validated this approach in humans with recurrent GBM. Combined, these data suggest an optimally administered dosage of 300 mg bid capecitabine, with higher doses yielding no significant gains in MDSC reduction while increasing the propensity for side effects and potential global immune suppression. Future studies should now investigate this approach in newly diagnosed GBM patients at the 300 mg bid dose, based on our observation that newly diagnosed patients have an overall increase in MDSC levels over time (which low-dose capecitabine was demonstrated to reduce, Figure 3A).

This study assessed the paradigm of an immunomodulatory approach based not on activating T cells but rather relieving immune suppression in the tumor microenvironment by targeting MDSCs. However, targeting MDSCs as a monotherapeutic strategy may not be sufficient to obtain a durable clinical response. Ongoing approaches are exploring immune-activating strategies in GBM, including immune checkpoint inhibitors, oncolytic viruses, CAR-T cells, and vaccine approaches. Unfortunately, none of these approaches have yet to demonstrate durable immune responses and convincing evidence of patient benefit^10, 44^. One possible explanation for this lack of clinical benefit is that the treatments tested have yet to overcome the inherent immune-suppressive nature of GBM. Therefore, future studies should be designed to utilize capecitabine in combination with other immune checkpoint inhibitors such as anti-PD1 therapy. This design is based on the observation that while CTLA-4 was reduced upon treatment, we also observed that PD-1 increased, possibly due to immune activation, in capecitabine-treated patients, making it a practical combinatorial treatment strategy to further enhance the anti-tumor immune response (**Supplemental Figure 5–7**). The immunobiology of GBM is complex, and given the negative results of other clinical immunotherapy trials in GBM, our findings suggest that any successful GBM immunotherapy strategy will likely require a synergistic approach that includes targeting of the immune suppressive/inhibitory effects of MDSCs along with utilization of immune activation strategies to overcome the global GBM-induced immunosuppression phenomenon.

## Methods

### Clinical study design

#### Clinical Trial Design

This trial is a phase 0/1 study of low-dose capecitabine and bevacizumab in patients with recurrent GBM (ClinicalTrials.gov #NCT02669173). Once GBM recurrence was identified on MRI, study-specific informed consent was obtained. Capecitabine was given 5-7 days prior to a clinically indicated surgical resection and then continued post-operatively in 28-day cycles with periodic blood draws to assess peripheral blood immune cell populations over the course of the trial (Figure 1). Patients were included if they had a recurrent histologically confirmed WHO grade 4 glioma for which a clinically indicated resection was planned. All subjects were at least 18 years of age, had no prior treatment with capecitabine or bevacizumab, and had a Karnofsky Performance status ≥ 60%. Patients were excluded if they were receiving other investigational agents or if they had a history of adverse reactions to compounds with similar chemical or biologic composition to capecitabine or bevacizumab, active infection with hepatitis B or C, or HIV, or other known malignancy within the past 2 years. To continue on the trial after resection was performed, patients were required to have histologically confirmed tumor recurrence (one patient with radiation necrosis was removed from the trial at this stage).

The treatment regimen began with capecitabine in the preoperative setting (5-7 days prior to surgery) with the final pre-operative dose the day of surgery. Post-operatively, capecitabine was resumed no sooner than 10 days after surgery after clearance from the surgical team and was given on days 1-28 in 28-day cycles. Bevacizumab was started on cycle 2 of capecitabine in the standard doses of 10 mg/kg IV on days 1 (± 3 days) and 15 (± 3 days) of each cycle. Magnetic resonance imaging was performed every 8 weeks. Treatment was continued until progressive disease according to RANO (Response Assessment in Neuro-oncology) criteria.

Dose escalation took place with the following doses of capecitabine: 150 mg bid (dose level 1), 300 mg bid (dose level 2), and 450 mg bid (dose level 3). As this trial was for patients with recurrent GBM, bevacizumab was included in addition to the low-dose capecitabine so that patients also received standard-of-care therapy (this combination at full doses has been proven to be well tolerated^45^).

The primary endpoint of the study was the degree of reduction in the concentration of circulating MDSCs. Secondary endpoints included concentration of tissue MDSCs and T-regulatory cells in resected GBM, safety and toxicity of continuous low-dose capecitabine alone and with standard dose bevacizumab, progression-free survival at 6 months rate (PFS_6_), progression free survival, and overall survival.

#### Specimen Collection Design

Peripheral blood pharmacodynamic (PBPD) assays were performed at the following timepoints: 1. Baseline (at trial enrollment), 2. Upon completion of low-dose oral capecitabine for 5-7 days pre-operatively, 3. Post-operative day 1, 4. Immediately before cycle 1 of post-operative capecitabine, 5. Immediately before the addition of bevacizumab (post-operative cycle 2), and 6. Every 4 weeks until patient removal from the trial. Tumor tissue was submitted for analysis at the time of surgery. See **Supplemental Table 2** for the full study calendar.

As a reference group for correlative studies, a secondary cohort of newly diagnosed GBM patients who were untreated prior to surgery was followed over time, with PBPD assays performed at similar time intervals including on the day of surgery, 2 weeks post-surgery, and every 2 months. These patients were the same as those previously analyzed and reported by our group^13^. To analyze changes in the tumor microenvironment that occur in response to the 7-day pre-operative capecitabine treatment, three newly diagnosed GBM tumor samples and one recurrent GBM tumor sample were obtained via the Cleveland Clinic Brain Tumor Tissue Bank.

### Study Assessment

#### Clinical Outcomes Analysis

Data collection and analysis were performed at Cleveland Clinic in May 2019. Demographic, clinical, and molecular pathology characteristics of each patient were obtained from the electronic health record and trial database. PFS was defined as the time from trial enrollment until the diagnosis of progression, and overall survival was defined as the time from initial histopathologic diagnosis until date of death.

#### Peripheral Blood Analysis

Peripheral blood analysis was performed at the Cleveland Clinic between 2017 and 2019, with flow cytometry performed as previously described^13^. In brief, peripheral blood mononuclear cells were isolated from whole blood via Ficoll gradient. All samples were processed less than twenty-four hours post-blood draw and then frozen in freezing medium for storage. Staining and analysis were performed using standard protocols previously described, with MDSCs marked by CD11b^+^, CD33^+^, and HLA-DR^−/lo^ and then further subdivided into granulocytic MDSCs (CD15^+^) and monocytic MDSCs (CD14^+^)^13, 24^. T regulatory cells were gated as CD3^+^, CD4^+^, CD25^+^, and CD127^−^, as previously described^13^. CD8^+^ T cells were gated on CD3^+^, CD8^+^, and CD4^−^. Concentrations of blood MDSCs, immune cells, and relevant secreted factors were measured at baseline, pre- and post-op, and after the addition of bevacizumab.

#### Tumor Tissue Analysis

Analysis of tumor tissue was performed cooperatively at the Cleveland Clinic and University of California Los Angeles (UCLA) in 2018. Tumor tissue samples were collected during surgery when extra tissue was safely available and was not obtained for all patients. Upon receiving tissue samples, tumors were digested in collagenase IV (STEMCELL Technologies) for 1 hour at 37°C before being mechanically dissociated via passage through a 40 µM filter. After dissociation, the single cell suspension was washed in cold RPMI medium before being counted and frozen in freezing medium for future use. Samples were then sent to UCLA, where CyTOF analysis was performed as previously described^13^, except analysis was performed only on the CD45+ fraction of the events collected. FCS files were normalized between runs using beads and the Nolan lab normalizer^46^. Analysis tools were used in R following the methods described by Nowicka et al.^47^

### Statistics

Baseline demographic and clinical characteristics were characterized by median and range for continuous variables and by frequency distribution and percentage of total for categorical variables. Software utilized for data processing and analysis included R Studio (Version 1.1.463, RStudio, Inc., Boston, MA), GraphPad PRISM (Version 6, GraphPad Software Inc., San Diego, CA), FlowJo (Version 10.5.0, FlowJo LLC, Ashland, OR), and JMP Pro (Version 14.0.0, SAS Institute Inc., Cary, NC). Statistical tests included two-sample Student’s t-tests for continuous variables, Pearson’s χ2 tests and Fisher’s exact test for categorical variables, and Kaplan-Meier curves for time-to-event analysis. Wilcoxon’s rank-sum test was used as the test of significance for T cell counts. A p value less than 0.05 was considered significant.

### Clinical study oversight

The clinical trial and retrospective study protocols were approved by the Institutional Review Board of Cleveland Clinic (#16-085) and the Protocol Review and Monitoring Committee (PRMC) of the Case Comprehensive Cancer Center (CASE7315). All participants provided written informed consent to participate in the clinical trial. The study was conceived, designed, initiated, and performed by the academic investigators. The authors confirm the accuracy and completeness of the data and analysis and the fidelity of the study to the protocol. All authors agreed to submit the manuscript for publication.

## Author Contributions

DMP, TJA, MMG, AGA, BO, MAV and JDL provided conceptualization and design; AMM and MAV performed the surgical collection of treated GBM tissue; TJA, AGA, DB, GR, MM, PH, MMG, and CAW performed the experiments; TJA, AGA, MMG, BO, DB, MAV, JDL and GR analyzed the data; DMP, TJA, MMG, BO, DB, MAV, JDL wrote the manuscript; HIK, MAV, and JDL provided financial support; and all authors provided final approval of the manuscript.

## Acknowledgements

We thank the patients treated at the Rose Ella Burkhardt Brain Tumor and Neuro-Oncology Center for donation of blood and tumor samples for this study, and we thank the staff of the Rose Ella Burkhardt Brain Tumor and Neuro-Oncology Center for their collaboration in acquiring samples. We thank the Case Comprehensive Cancer Center and the Musella Foundation for their support of this clinical trial. We thank the members of the Lathia laboratory for insightful discussion and constructive comments on the manuscript. We thank Joseph Gerow and Eric Schultz for flow cytometry assistance. We thank the Janis V. Giorgi Flow Cytometry Core Laboratory (JCCC; UCLA) for their assistance with CyTOF experiments. We thank Amanda Mendelsohn and the Center for Medical Art and Photography at the Cleveland Clinic for providing illustrations and Dr. Erin Mulkearns-Hubert for editorial assistance. This work was funded by an NIH grant (F31 NS101771 to TJA), Cancer Biology Training Grant (T32CA059366 to DB), the Sontag Foundation (JDL), Blast GBM (JDL, MAV), the Cleveland Clinic VeloSano Bike Race (JDL, MAV), B*CURED (JDL, MAV), and the Case Comprehensive Cancer Center (JDL, MAV). Flow cytometry was performed in the UCLA JCCC and Center for AIDS Research Flow Cytometry Core Facility, which is supported by NIH awards P30 CA016042 and 5P30 AI028697. The purchase of the Helios mass cytometer that was used in this work was supported, in part, by the funds provided by the James B. Pendleton Charitable Trust. HIK and AGA were supported by the Dr. Miriam and Sheldon G. Adelson Medical Research Foundation.

