## Supplemental Figures for "Metronomic capecitabine as an immune modulator in glioblastoma patients reduces myeloid-derived suppressor cells"

**Supplemental Table 1:** Clinical Characteristics of study patients

| | Age at Diagnosis | Age at Study Enrollment | Sex | KPS at Diagnosis | Location | Prior Immunotherapy | Prior Chemotherapy | Initial Surgery | Recurrence # | Study Surgery | Study Dose | Progression at Time of Publication | Alive at the Time of Publication | Progression Free Survival | Overall Survival | On Cycle # when stopped | 1p Status | 19q Status | IDH Mutation Status | MGMT Methylation Status | Ki 67 Percent | EGFR Amplification | P53 Percent Reactivity Percent |
|---|---|---|---|---|---|---|---|---|---|---|---|---|---|---|---|---|---|---|---|---|---|---|---|
| 1 | 56 | 57 | M | 90 | Right frontal | No | Yes | Biopsy | 1 | STR | 300 | Yes | No | 5.5 | 17.5 | 5 | Loss | Intact | Wild Type | Unmethylated | 40 | No | 10 |
| 2 | 64 | 65 | M | 80 | Left parietal | No | No | NTR | 1 | NTR | 150 | Yes | No | 9.2 | 14.1 | 10 | Loss | Loss | Wild Type | Unmethylated | 70 | Yes | 25 |
| 3 | 67 | 70 | F | 90 | Left occipital | Yes | No | NTR | 2 | NTR | 300 | Yes | No | 1.8 | 29.7 | 2 | Intact | Intact | Wild Type | Methylated | 50 | Yes | 25 |
| 4 | 63 | 64 | F | 80 | Left frontal | Yes | No | GTR | 1 | NTR | 150 | Yes | No | 5.3 | 35.4 | 5 | Intact | Intact | Wild Type | Unmethylated | 50 | Yes | 10 |
| 5 | 38 | 40 | F | 90 | Right frontal | No | Yes | GTR | 2 | GTR | 150 | No | Yes | 27.8 | 36.6 | 30 | NA | NA | Mutated | Methylated | 12 | NA | 75 |
| 6 | 58 | 59 | M | 90 | Right temporal | No | Yes | Biopsy | 2 | NTR | 300 | Yes | No | 5.8 | 22.1 | 6 | Intact | Intact | Wild Type | Unmethylated | 40 | Yes | 5 |
| 7 | 58 | 59 | M | 80 | Right frontal | No | No | NTR | 1 | GTR | 150 | Yes | No | 5.1 | 14.9 | 4 | NA | NA | Wild Type | Methylated | 30 | NA | NA |
| 8 | Removed from trial after surgical pathology results showed no disease progression | | | | | | | | | | | | | | | | | | | | | | |
| 9 | 56 | 58 | M | 90 | Right parietal | No | No | GTR | 1 | GTR | 450 | Yes | No | 1.8 | 11.9 | 2 | Intact | Intact | Wild Type | Unmethylated | 70 | Yes | 10 |
| 10 | 60 | 61 | F | 80 | Left frontal | Yes | Yes | GTR | 2 | NTR | 450 | Yes | No | 8.5 | 21.6 | 9 | Intact | Intact | Wild Type | Methylated | 9 | No | 10 |
| 11 | 45 | 46 | F | 90 | Left Parietal | No | No | GTR | 1 | STR | 450 | No | Yes | 13.9 | 22.3 | 14 | Intact | Loss | Wild Type | Methylated | 80 | No | 5 |
| 12 | 48 | 49 | M | 90 | Right temporal | No | No | Laser | 1 | GTR | 450 | Yes | Yes | 6.2 | 12.9 | 7 | Intact | Intact | Wild Type | Unmethylated | 40 | No | 20 |

Abbreviations: M: male; F: female; KPS: Karnofsky Performance Scale; STR: subtotal resection; NTR: near-total resection; GTR: gross-total resection; IDH: isocitrate dehydrogenase; MGMT: O-6-methylguanine-DNA methyltransferase; EGFR: epidermal growth factor receptor

**Supplemental Table 2:** Study Calendar

| Required Assessments | Pre- study | Pre-op | Post op (Day 1) | Cycle 1 (Day 1)[1] | Cycle 2+ (Day 1)[1] | End of Treatment | 30-Day Follow up[10] |
|---|---|---|---|---|---|---|---|
| Informed Consent | X | | | | | | |
| Demographics | X | | | | | | |
| Medical History | X | | | X | X | X | X |
| Weight | X | X[11] | | X | X | X | X |
| Vitals | X | X[11] | | X | X | X | X |
| Physical Exam | X | | | X | X | X | X |
| Con Meds | X | | | X | X | X | X |
| KPS | X | | | X | X | X | X |
| Baseline Symptoms | X | | | | | | |
| AE Assessment | X | X[11] | | X | X | X | X |
| CBC/diff | X | | | X | X | X | |
| CMP | X | | | X | X | X | |
| Urine P/C ratio | X | | | | X[9] | | |
| PT/INR | X | | | | | | |
| Pregnancy test[8] | X | | | | | | |
| ECG | X | | | | | | |
| MRI[7] | X | | | | X | X | |
| Capecitabine | | X[3] | | X | X | | |
| Bevacizumab | | | | | X[2] | | |
| Correlative Studies | | X[4,5,12] | X | X[4] | X[4,6] | | |

1 – ± 3 days
2 – Days 1 and 15 (± 3 days)
3 – Capecitabine to start 5-7 days before surgery with last dose on morning of surgery
4 – Perform up to 3 days before starting chemotherapy
5 – Tissue studies on resected specimen
6 – Repeat through Cycle 6 and thereafter at the discretion of the investigator
7 – MRI to be done with gadolinium. Repeat MRI before (up to 5 days) even cycles beginning with Cycle 2. Perfusion MRI scanning will be done according to the discretion of the treating investigator.
8 – Only if not done pre-op and if child-bearing potential; serum or urine test allowed
9 – Omit test on Cycle 2, Day 1
10 – ± 2 weeks
11 – May be combined with Pre-study assessments
12 – Blood sample preoperatively on day of surgery

**Supplemental Figure 1**

A

| MDSC panel | | |
|---|---|---|
| **Ab** | Fluorophore | cat # |
| **CD11b** | AF700 | CD11b29 |
| **HLA-DR** | APC | 559866 |
| **CD14** | APC H7 | 560180 |
| **CD15** | PerCP | 555400 |
| **CD33** | PE | 555450 |

| T cell panel | | |
|---|---|---|
| **Ab** | Fluorophore | Cat # |
| **CD4** | PerPC | 347324 |
| **CD3** | 488 | 555332 |
| **CD25** | PE | 555432 |
| **CD8** | PE cy7 | 557746 |
| **CD107a** | APC-H7 | 561343 |
| **Cd127** | Af647 | 558598 |

B

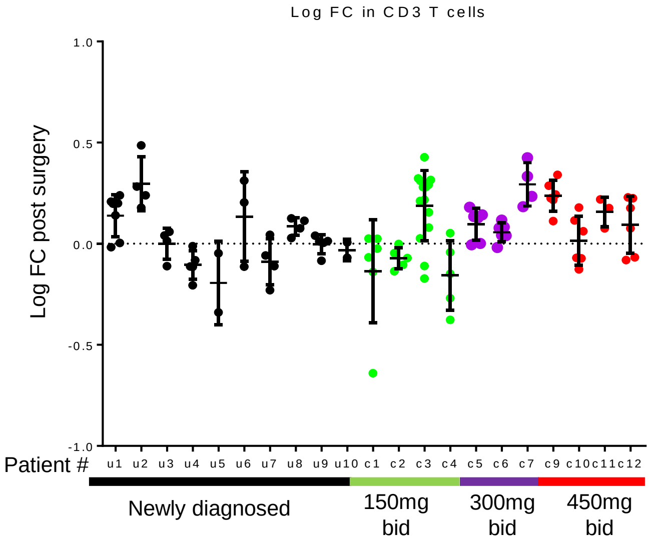


C

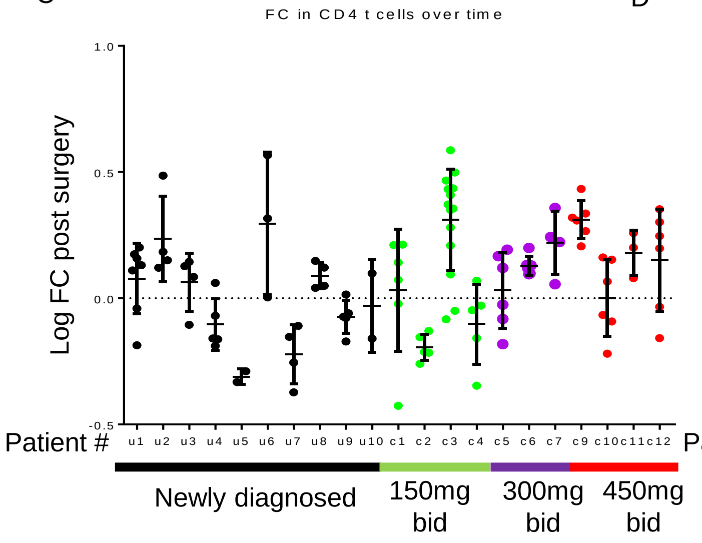


D

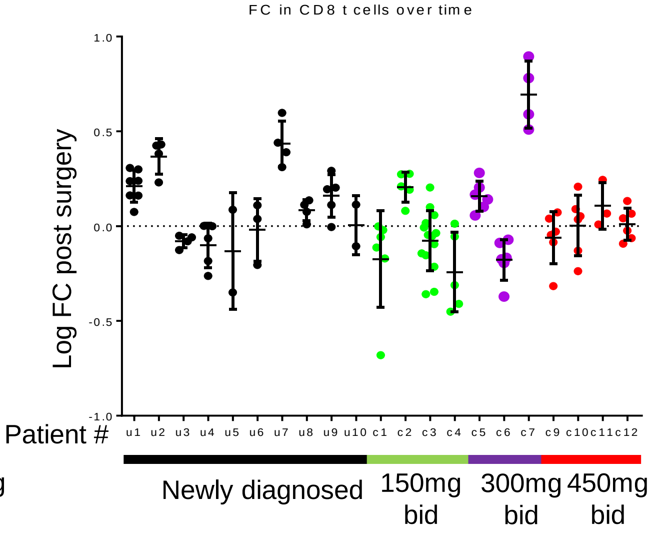


E

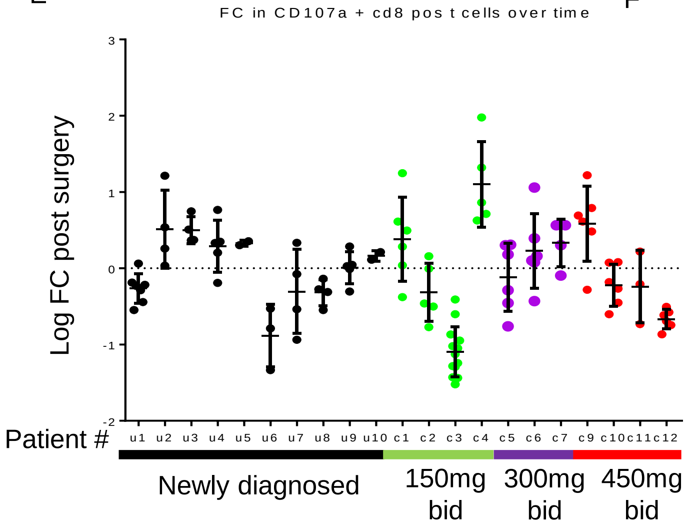


F

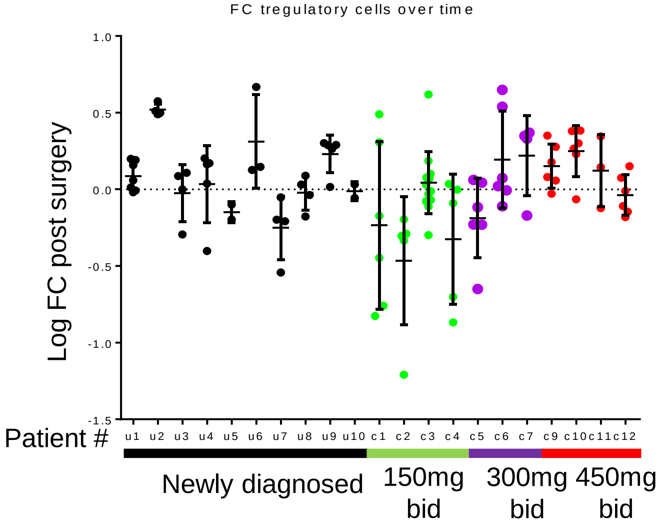

### Supplemental Figure 2

A

| label | target | Clone # | Cat # |
|---|---|---|---|
| 209Bi | CD11b (Mac-1) | ICRF44 | 3209003B |
| 170Er | CD3 | UCHT1 | 3170001B |
| 167Er | CD197 (CCR7) | G043H7 | 3167009A |
| 165Ho | CD61 | VI-PL2 | 3165010B |
| 164Dy | CD15 (SSEA-1) | W6D3 | 3164001B |
| 163Dy | CD56 (NCAM) | NCAM16.2 | 3163007B |
| 146Nd | CD8a | RPA-T8 | 3146001B |
| 159Tb | CD11c | Bu15 | 3159001B |
| 158Gd | CD33 | WM53 | 3158001B |
| 169Tm | CD45RA | HI100 | 3169008B |
| 89Y | CD45 | HI30 | 3089003B |
| 153Eu | TIM-3 | F38-2E2 | 3153008B |
| 151Eu | CD123 (IL-3R) | 6H6 | 3151001B |
| 150Nd | CD223 (LAG-3) | 11C3C65 | 3150030B |
| 149Sm | CD66a | CD66a-B1.1 | 3149008B |
| 148Nd | CD16 | 3G8 | 3148004B |
| 147Sm | CD20 | 2H7 | 3147001B |
| 145Nd | CD4 | RPA-T4 | 3145001B |
| 143Nd | CD25 (IL-2R) | M-A251 | 555430 |
| 142Nd | CD19 | HIB19 | 3142001B |
| 139La | CD107a (LAMP1) | H4A3 | 328635 |
| 174Yb | HLA-DR | L243 | 3174001B |
| 155Gd | CD279 (PD-1) | EH12.2H7 | 3155009B |
| 176Yb | CD127 (IL-7Ra) | A019D5 | 3176004B |
| 160Gd | CD28 | CD28.2 | 3160003B |
| 161Dy | CD152 (CTLA-4) | 14D3 | 3161004B |
| 175Lu | CD14 | M5E2 | 3175015B |
| 171Yb | CD68 | Y1/82A | 3171011B |

B

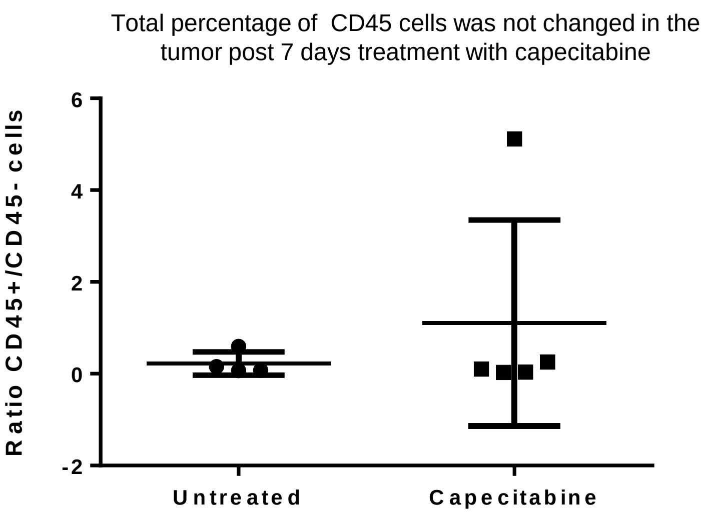


C

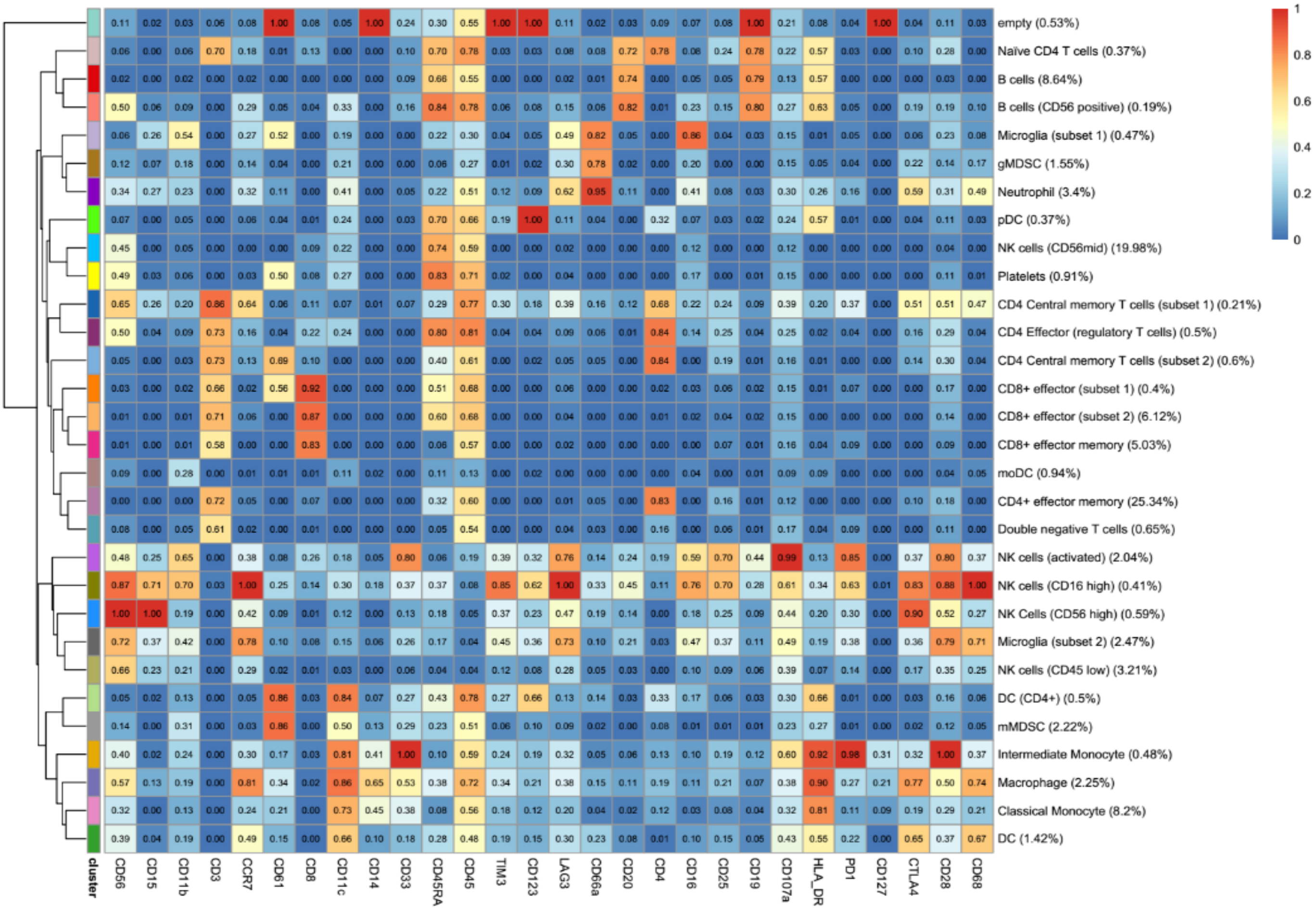

**Supplemental Figure 3**

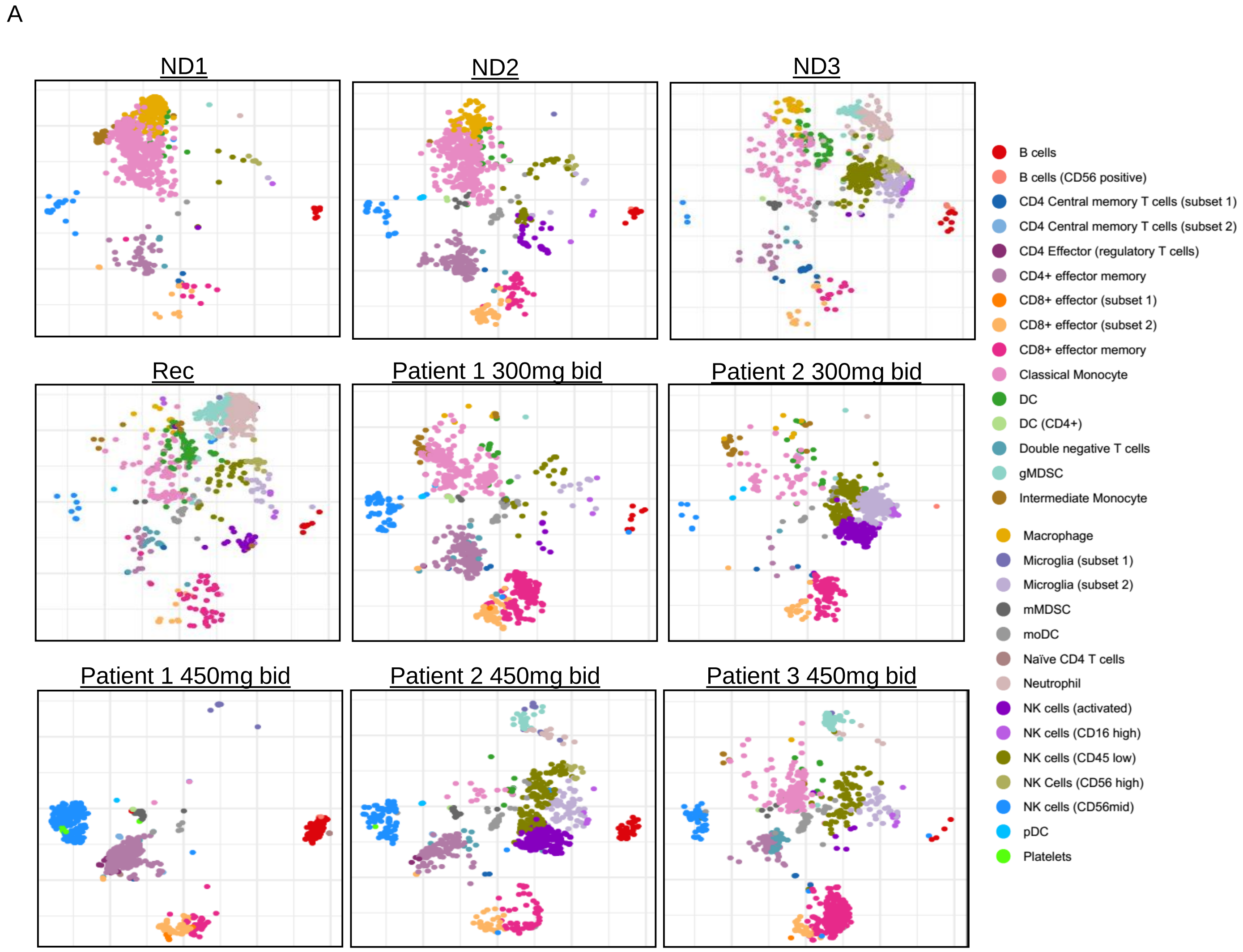

**Supplemental Figure 4**

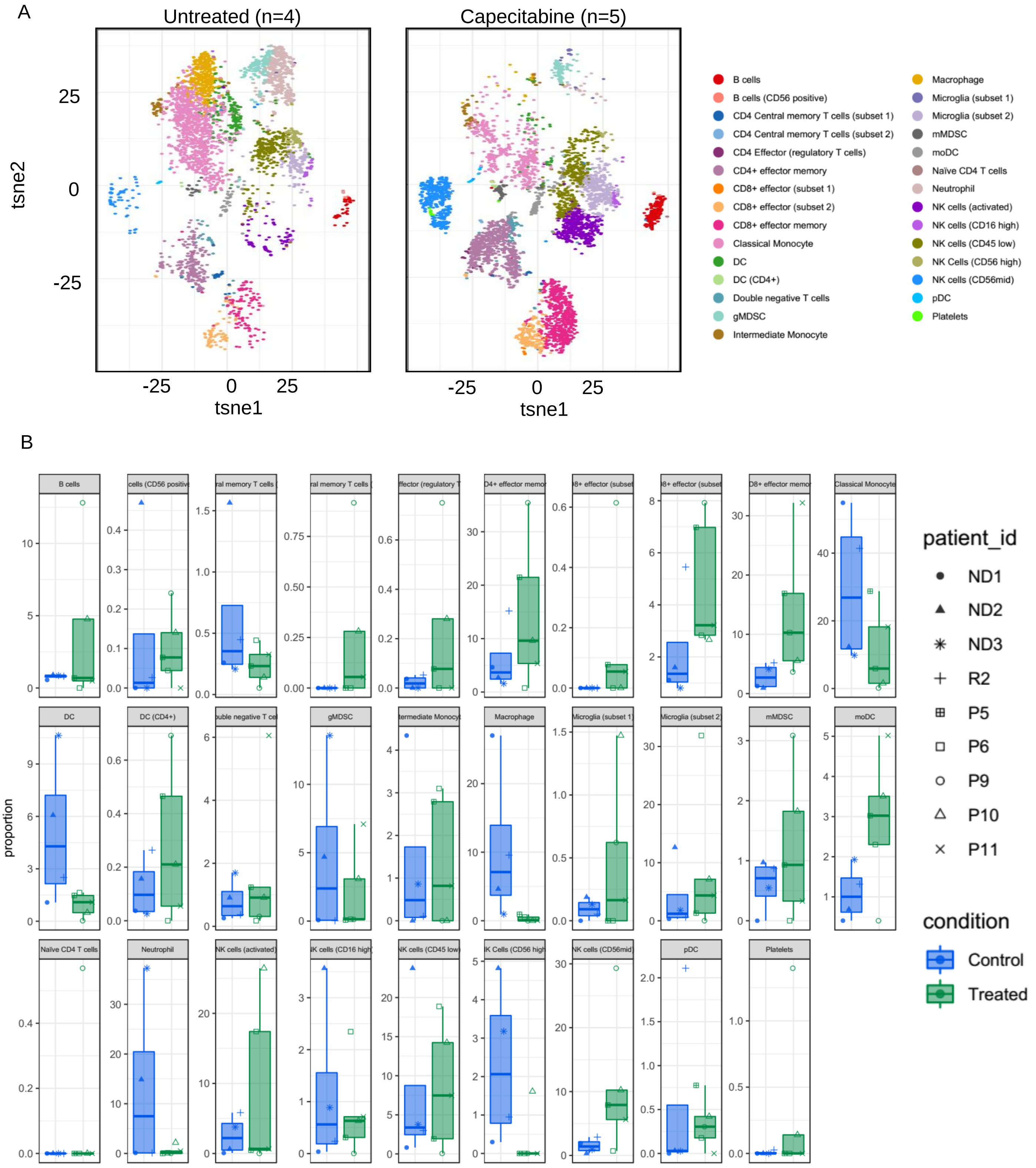

**Supplemental Figure 5**

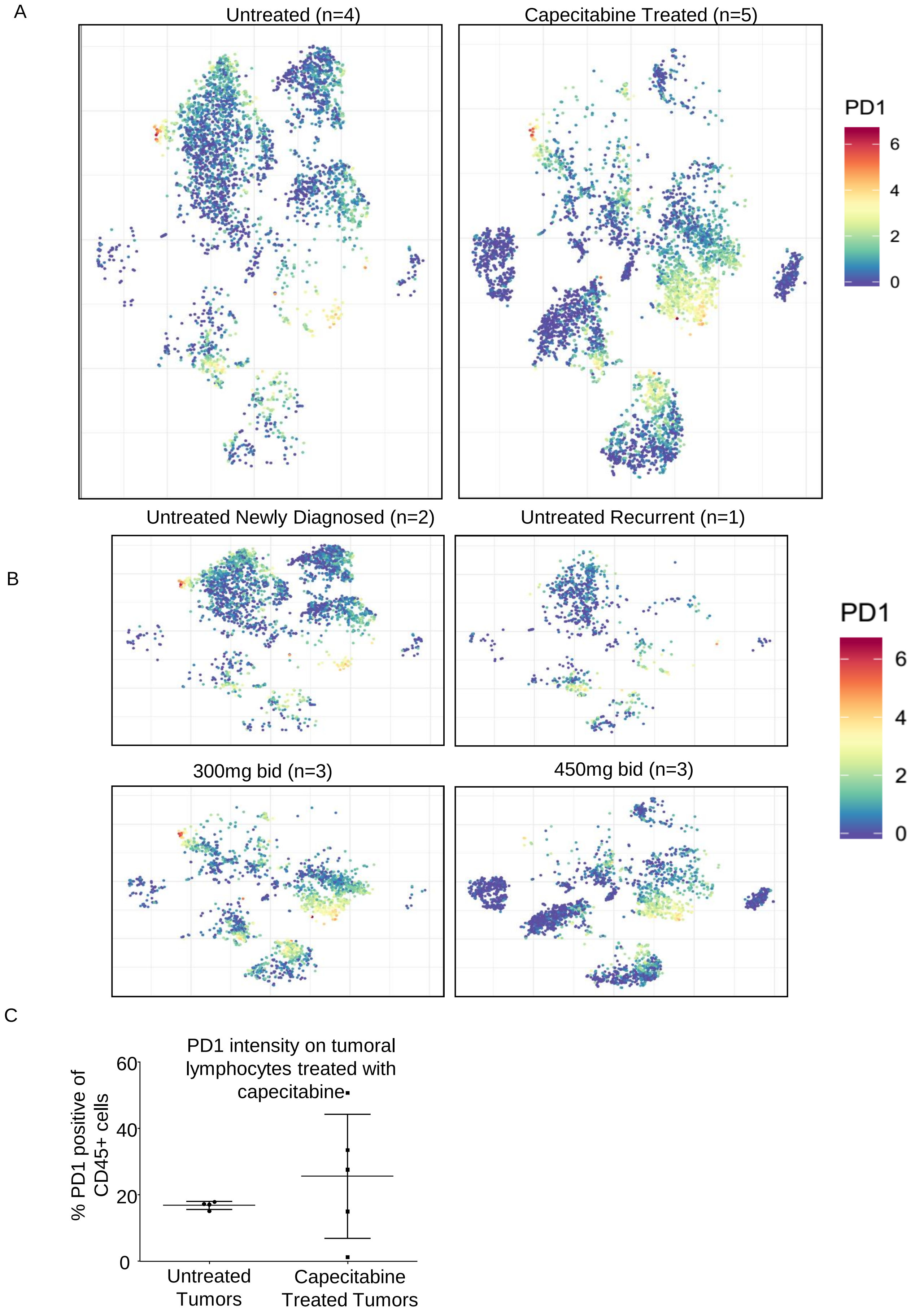

**Supplemental Figure 6**

A

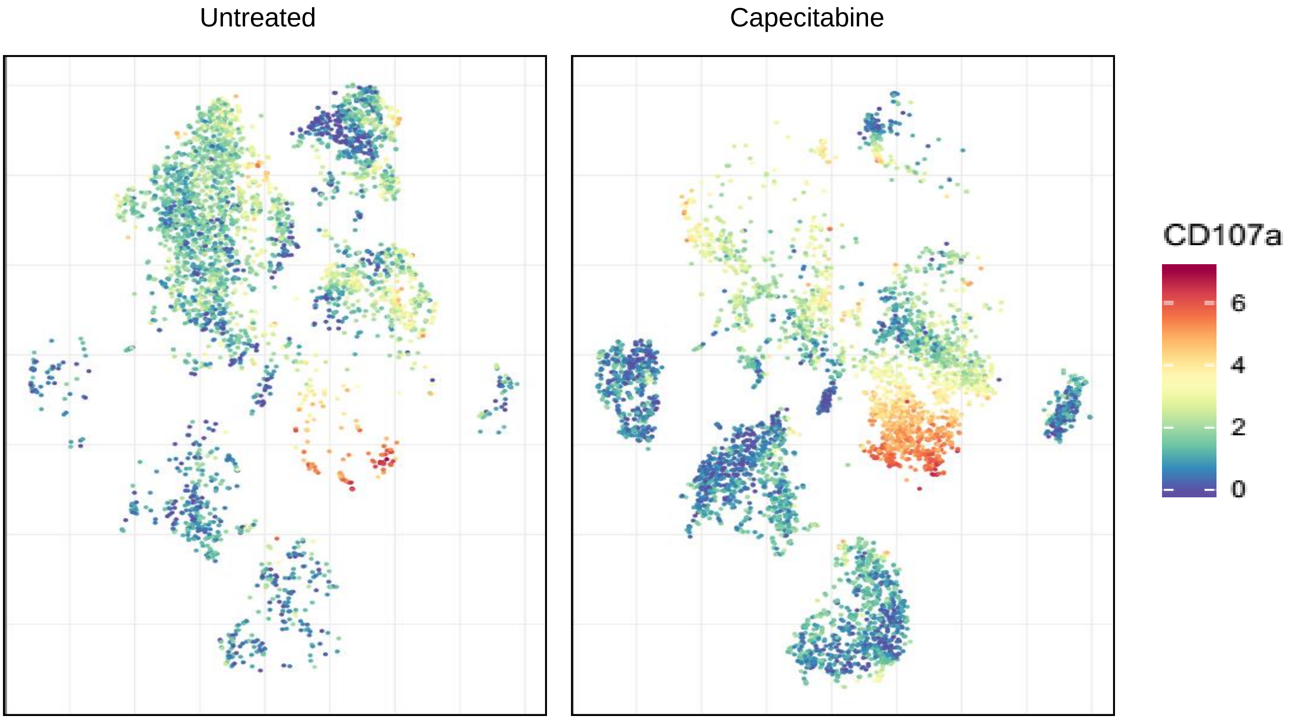


B

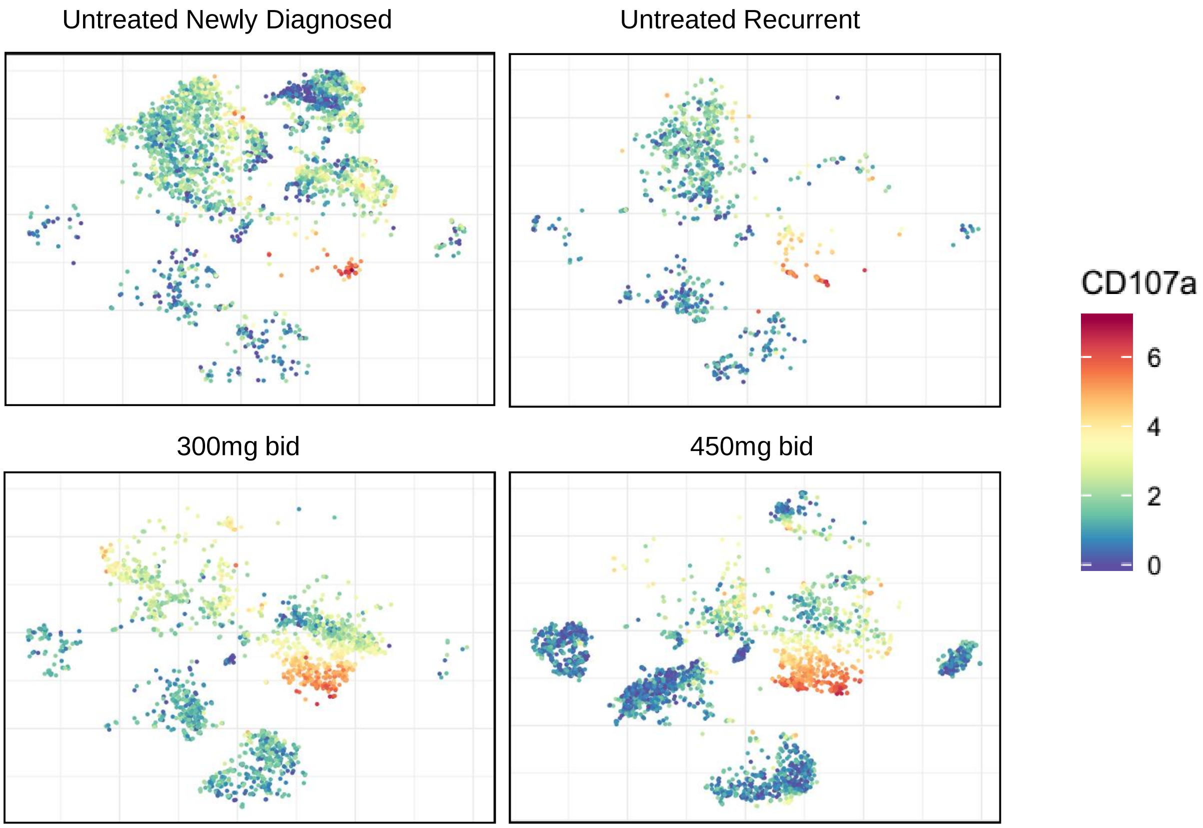

**Supplemental Figure 7**

A

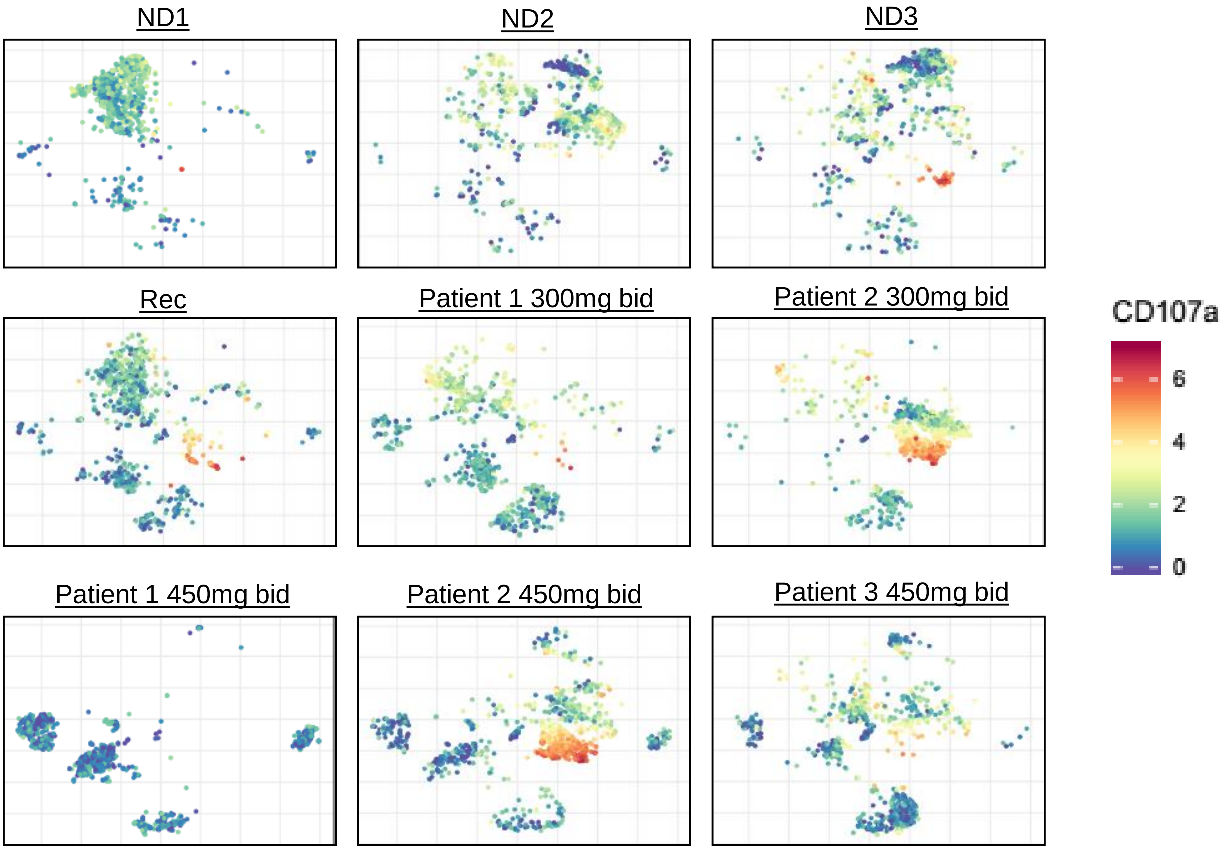


B

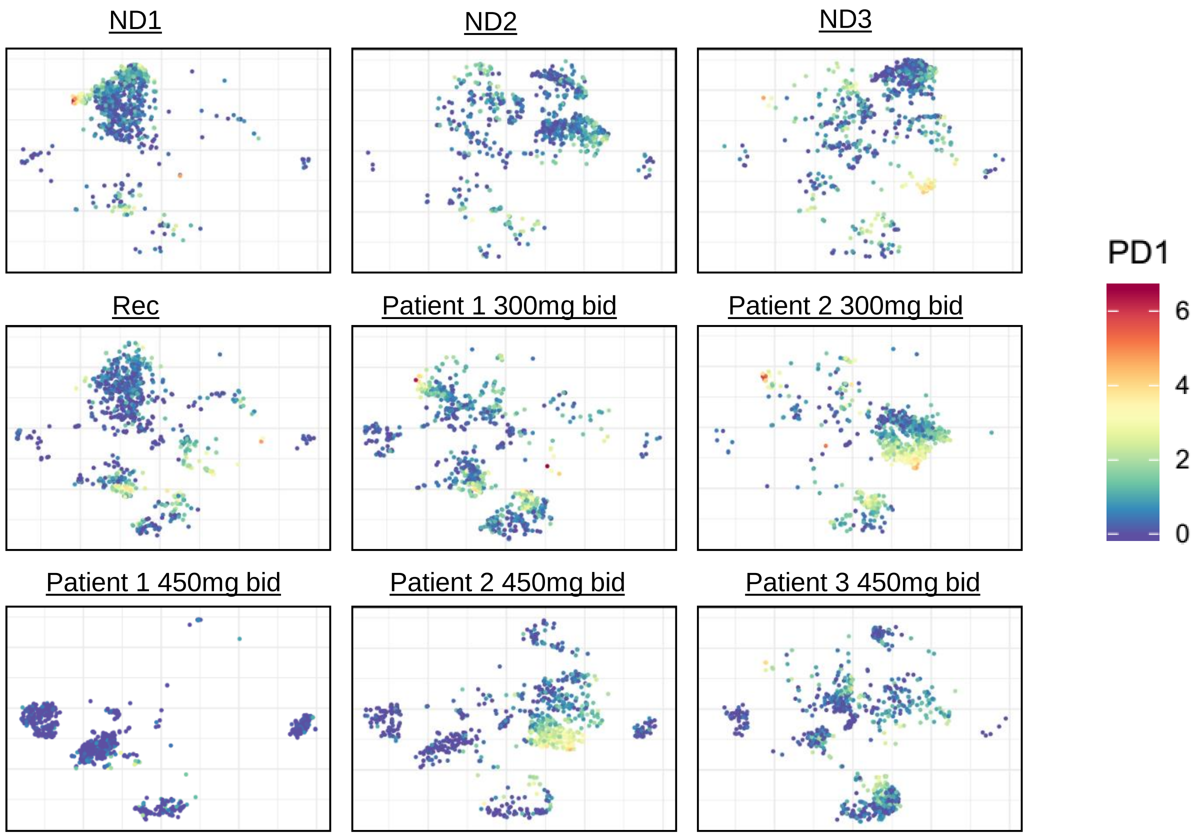

**Supplemental Table Descriptions**

**Supplemental Table 1.** Detailed characteristics, including demographics, clinical and treatment parameters, and tumor-specific markers, of the 11 evaluable patients.

**Supplemental Table 2.** Study calendar detailing the required assessments and their respective acquisition time points

**Supplemental Figure Legends**

**Supplemental Figure 1.** Post-surgical resection, the circulating T cell populations did not change compared to controls in response to capecitabine treatment. Flow cytometry was performed on PBMC samples collected using two flow cytometry panels (MDSC panel, and T cell panel) (**A**). Log fold change in total CD3+ T cells from surgical resection over time showing a newly diagnosed cohort of patients not treated with capecitabine (**B**). ($CD3^+$, $CD4^+$) CD4+ T cells, ($CD3^+$, $CD8^+$) T cells, ($CD3^+$, $CD8^+$, $CD107a^+$) cytotoxic T cells, and ($CD3^+$, $CD4^+$, $CD127^-$, $CD25^+$) T regulatory cells were also analyzed over time in each patient, demonstrating no trend upon capecitabine treatment in any of these lymphocyte populations (**C-F**).

**Supplemental Figure 2.** CyTOF immune panel analysis. CyTOF analysis utilized an immune panel consisting of the immune markers listed in (**A**). The CyTOF was performed on whole dissociated tumor samples, with an initial step of separating CD45+ cells from CD45- cells. The total number of CD45+ cells was not different between the untreated and capecitabine-treated patients (**B**). Unbiased clustering was used to identify the following immune populations by heatmap analysis for each marker of the panel (**C**). All error bars represent the standard deviation. Unpaired student's t-test was used for all comparisons, where *$p < 0.05$, **$p < 0.01$, ***$p < 0.001$.

**Supplemental Figure 3.** CyTOF comparison of capecitabine-treated vs untreated patients. Comparison of treated (n=5) vs control samples (n=4) samples represented as a tSNE plot (**A**). The various immune population names are denoted to the right in the figure key. Graphical representation of treated vs untreated patients with each of the 29 immune populations identified (**B**).

**Supplemental Figure 4.** Multidimensional plots of immune populations from individual patients (**A**). Individual population color key is located to the right of the figures.

**Supplemental Figure 5.** PD-1 levels of untreated and treated patients were compared using tSNE multidimensional plots, with PD-1 expression levels colored according to the key to the right of the figure (**A**). Subdividing the tSNE plots of treated and untreated patients for PD-1 levels of newly diagnosed patients, recurrent patients, 300 mg bid capecitabine, and 450 mg bid capecitabine (**B**). Quantification of PD-1+ cells of the CD45+ lymphocytes did not identify any significant differences between untreated and treated tumor samples.

**Supplemental Figure 6.** CD107a levels of untreated and treated patients were compared using tSNE multidimensional plots, with CD107a expression levels colored according to the key to the right of the figure (**A**). Subdividing the tSNE plots of treated and untreated patients for CD107a levels for newly diagnosed patients, recurrent patient, 300 mg bid capecitabine, and 450 mg bid capecitabine (**B**).

**Supplemental Figure 7.** CD107a and PD-1 levels are depicted using individual patient-based tSNE plots (**A, B**).
